# Macroecological dynamics of gut microbiota

**DOI:** 10.1101/370676

**Authors:** Brian W. Ji, Ravi U. Sheth, Purushottam D. Dixit, Konstantine Tchourine, Dennis Vitkup

## Abstract

The gut microbiome is now widely recognized as a dynamic ecosystem that plays an important role in health and disease^1^. While current sequencing technologies make it possible to estimate relative abundances of host-associated bacteria over time^2, 3^, the biological processes governing their dynamics remain poorly understood. Therefore, as in other ecological systems^4, 5^, it is important to identify quantitative relationships describing global aspects of gut microbiota dynamics. Here we use multiple high-resolution time series data obtained from humans and mice^6–8^ to demonstrate that despite their inherent complexity, gut microbiota dynamics can be characterized by several robust scaling relationships. Interestingly, these patterns are highly similar to those previously observed across diverse ecological communities and economic systems, including the temporal fluctuations of animal and plant populations^9–12^ and the performance of publicly traded companies^13^. Specifically, we find power law relationships describing short- and long-term changes in gut microbiota abundances, species residence and return times, and the connection between the mean and variance of species abundances. The observed scaling relationships are altered in mice receiving different diets and affected by context-specific perturbations in humans. We use these macroecological relationships to reveal specific bacterial taxa whose dynamics are significantly affected by dietary and environmental changes. Overall, our results suggest that a quantitative macroecological framework will be important for characterizing and understanding complex dynamics of microbial communities.

---

The dynamics of gut bacteria can now be monitored with high temporal resolution using 16S rRNA amplicon sequencing^14^. Recent longitudinal studies have revealed significant day to day variability and marked long-term stability of gut microbiota^6, 7, 15, 16^. Several studies have also identified important factors, such as host diet and lifestyle, that contribute to temporal changes in species abundances^7, 8, 17, 18^. However, in contrast to other macroscopic ecological communities, statistical relationships describing gut microbiota dynamics are not well understood. While ideas from theoretical ecology have been applied to understand static patterns of gut microbial diversity and species abundance distributions^19, 20^, a comprehensive, quantitative analysis of macroecological dynamics is currently missing. Therefore, we sought to investigate dynamical relationships in the gut microbiome using several of the longest and most densely-sampled longitudinal studies in humans and mice^6–8^. The considered data spanned three independent investigations, utilizing different sample collection procedures and sequencing protocols; bacterial abundances in these studies were tracked daily for several weeks in mice and up to a year in humans. Our analysis included four healthy human individuals (A, B, M3, F4) and six individually-housed mice fed either a low-fat, plant polysaccharide (LFPP) diet or a high-fat, high-sugar (HFHS) diet. We use these data to explore the short-term abundance changes and long-term drift of gut microbiota, species residence and return times, and the temporal variability of individual bacterial taxa across humans and different mouse diet groups. Collectively, our study provides a comprehensive characterization of macroecological dynamics in the gut microbiome.

Following a quantitative framework used previously to examine the ecological dynamics of animal populations^9, 10^, we first investigated short-term temporal fluctuations of gut microbiota abundances. One of the most basic descriptors of bacterial population dynamics is the daily abundance change, defined as the logarithm of the ratio of consecutive daily abundances, *μ*_*k*_(*t*) = log(*X*_*k*_(*t* + 1)/*X*_*k*_(*t*)), where *X*_*k*_ is the relative abundance of a bacterial operational taxonomic unit (OTU) *k* at time *t*. Defined in this way, *μ*(*t*) quantifies the rate of change of OTU abundances averaged over the course of a day. Interestingly, we found that the probability of *μ* averaged over all OTUs closely followed a Laplace distribution (Equation 1), with a characteristic tent shape in log-transformed probabilities (Fig. 1a-c).

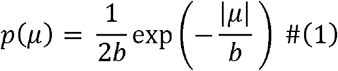

**Fig. 1.**
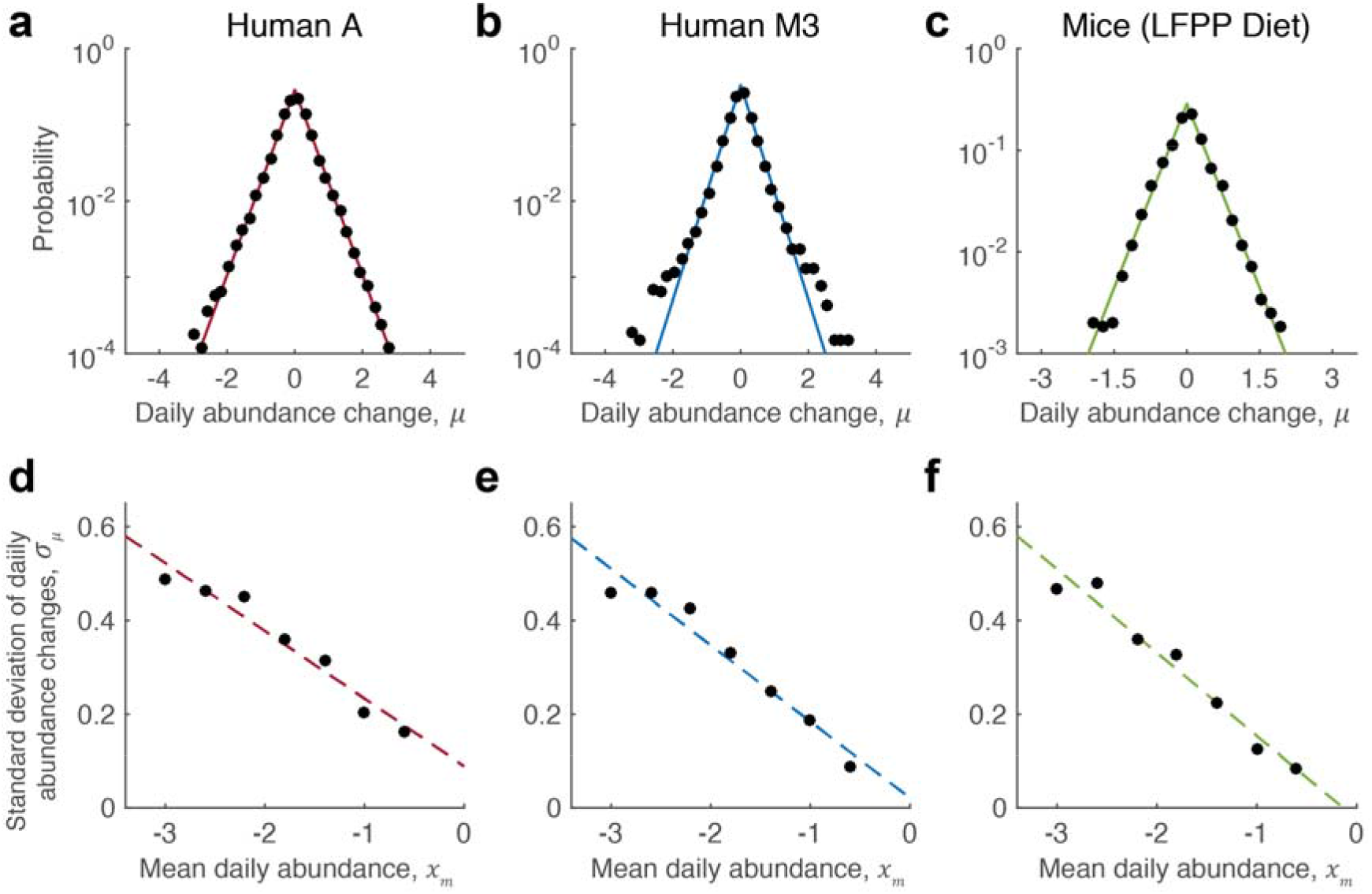
Daily changes in the abundances of gut microbiota. **a-c,** Daily abundance changes were defined as *μ*_*k*_(*t*) = log(*X*_*k*_(*t* + 1)/*X*_*k*_(*t*)), where *X*_*k*_ is the relative abundance of a given OTU *k* on day *t*. The distribution of *μ* averaged over all OTUs displays a Laplace form (Equation 1), appearing as a characteristic tent shape in log-transformed probabilities. Results are shown for two individuals from different human studies (A and M3) and mice fed a low-fat plant polysaccharide-based (LFPP) diet. Laplace exponents are *b* = 0.83 ± 0.1 for human A, *b* = 0.71 ± 0.07 for human M3, and *b* = 0.82 ± 0.10 for LFPP mice (mean ± s.d., Methods). Solid lines indicate fits to the data using maximum likelihood estimation (MLE). **d-f,** Across all OTUs, the standard deviation of daily abundance changes (*σ*_*μ*_) decreases with mean daily abundance (*x*_*m*_), defined as the mean of successive log abundances, 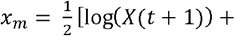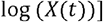. Standard deviations were calculated by binning daily abundance changes by different values of *x*_*m*_ along the x-axis. Dashed lines are least-squares fits to the data, with slopes of *r* = −0.16 ± 0.02, −0.16 ± 0.02 and −0.17 ± 0.03 for A, M3 and LFPP mice respectively (mean ± s.d., see Methods). Abundance changes in (**c**) and (**f**) were aggregated across the three mice on the LFPP diet.

Laplace distributions were highly similar within and between individual humans, and between humans and mice (parameter *b* = 0.73 ± 0.07, *b* = 0.82 ± 0.1; mean ± s.d. across all humans and LFPP mice respectively), indicating the universality of these relationships. Moreover, the Laplace distribution described well the daily abundance changes of every gut microbiome time series we analyzed (Supplementary Fig. 1), including those defined at various taxonomic resolutions (Supplementary Fig. 1c). We note that the observed distributions are unlikely to arise due to species aggregation (Supplementary Fig. 1d)^21, 22^ or compositional nature of bacterial abundance data we use (Supplementary Fig. 3).

In contrast to the Gaussian distribution (see Supplementary Figs. 1 and 2 for model fits), which is expected when bacterial growth is affected by random multiplicative processes^20, 23^, the Laplace distribution indicates substantially higher probabilities for large short-term bacterial abundance fluctuations. A Laplace distribution of abundance variability may arise due to density-dependent birth and death rates in a migrating population^12^ or through emergence of sub-specialized environmental niches^24^. Nevertheless, the exact mechanisms and dynamic processes generating this distribution are currently not well-understood and need to be investigated further. The symmetry of the Laplace distribution suggests an equal probability for increases or decreases in species’ abundances, which reflects a zero-sum process due to finite resources in the gut. Interestingly, Laplace distributions have been observed across many diverse ecological and economic systems including bird communities^9, 10^, fish populations^11^, tropical rain forests^12^, publicly traded company sales^13^, and country GDPs^25^ (Supplementary Fig. 4a). Similar to these complex ecological and interacting systems, the gut microbiome may often exhibit sudden large-scale abundance fluctuations.

In many complex ecosystems, species short-term abundance fluctuations often depend on their current abundance^9, 10, 13, 25^. We therefore investigated the relationship between the species’ abundances and the standard deviation of daily abundance changes. The analysis revealed that the variability of daily abundance changes of gut bacteria decreased approximately linearly with increasing mean daily abundances (Fig. 1d-f). This result was not due to sampling errors associated with finite sequencing depth (Supplementary Fig. 5), and the decrease in daily abundance changes was also observed at the single OTU level (Supplementary Fig. 6). Moreover, the observed behavior was similar between human and mouse gut microbiomes (regression slopes *r* = −0.15± 0.01, −0.17± 0.03; mean ± s.d. across humans and mice). Thus, likely due to the presence of more stable nutrient niches, highly abundant bacteria exhibit substantially smaller relative daily fluctuations compared to bacteria with lower abundances.

In addition to short-term dynamics, interesting long-term dynamical trends have also been observed across different macroscopic ecosystems^9, 21, 26^. To explore the long-term behavior of gut microbiota, we investigated how the mean-squared displacement (MSD) of OTU abundances (⟨*δ*^2^(Δ*t*)⟩) changed with time. Again, similar to the behavior of other diverse communities (Supplementary Fig. 4c), we found that the long-term dynamics of gut microbiota abundances could be well approximated by the equation of anomalous diffusion (Fig. 2, Supplementary Fig. 7),

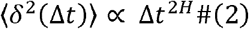

where *H* is the Hurst exponent quantifying the collective rate of abundance drift over time and therefore, the long-term stability of gut microbiota^27^. In comparison with normal diffusion (*H* = 0.5), a Hurst exponent of *H* > 0.5 indicates a tendency for increases (decreases) in abundances to be followed by further increases (decreases), whereas a value of *H* < 0.5 indicates a higher degree of stability and a bias for abundances to revert back to their means. In contrast to short-term fluctuations of bacteria abundances, described by the Laplace distribution (Equation 1, Fig. 1), the Hurst exponent in Equation 2 quantifies the rate at which the average root mean squared displacement of abundances increases as a function of time. Both in human and mouse gut microbiomes, our analysis revealed small Hurst exponents (*H* = 0.09 ± 0.03, *H* = 0.08 ± 0.02, mean ± s.d. across humans and mice). This suggests that despite overall stability^15, 28, 29^, gut microbiota exhibit a slow but continuous and predictable long-term abundance drift. Furthermore, while the temporal behavior of individual OTU abundances was also well-approximated by the equation of anomalous diffusion (Supplementary Fig. 8a), the distribution of Hurst exponents across individual OTUs exhibited substantial variability (Supplementary Fig. 8b). This demonstrates the heterogeneity in the stability of different gut bacterial taxa within and across hosts. We show below that the stability of different taxa can be significantly affected by environmental factors such as host dietary intake.

**Fig. 2.**
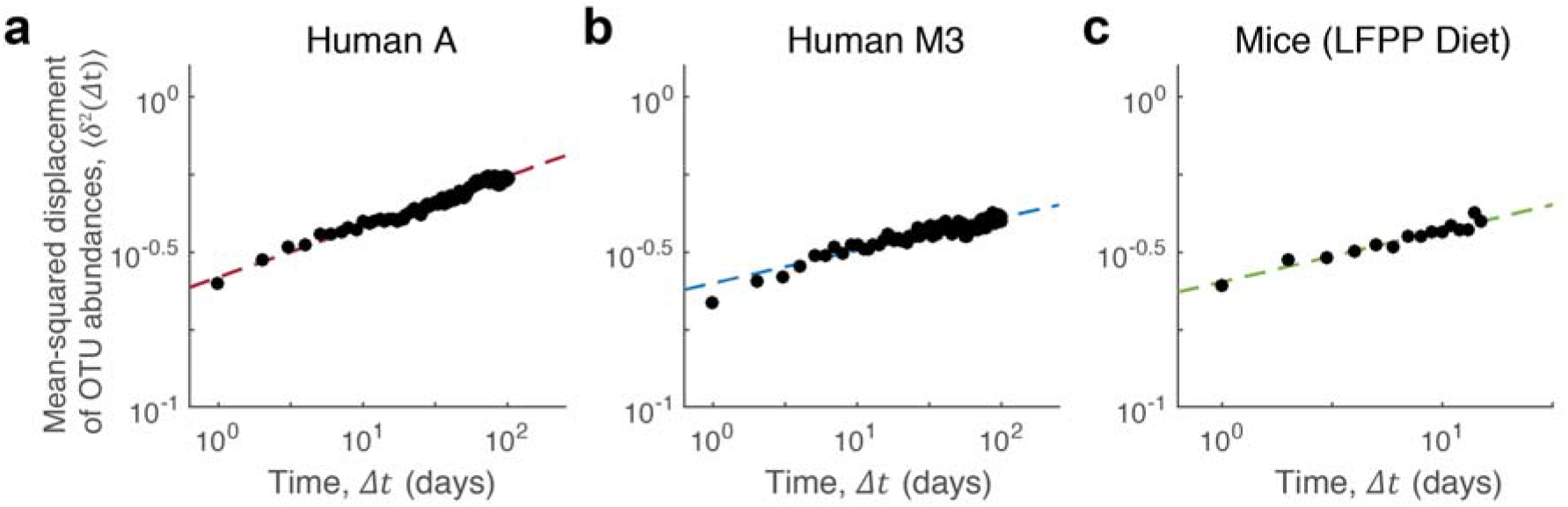
Long-term stability of gut microbiota abundances. **a-c,** In humans and mice, the mean-squared displacement of log OTU abundances (⟨*δ*^2^(Δ*t*)⟩) scales with time as a power law of the form ⟨*δ*^2^(Δ*t*)⟩ ∝ Δ*t*^2*H*^. Hurst exponents are *H* = 0.07 ± 0.03, 0.08 ± 0.02, 0.08 ± 0.02 for human A, human M3 and LFPP mice respectively (mean ± s.d., Methods). The data in (**c**) represent an average over the three individual mice on the LFPP diet (Methods). Dashed lines indicate least-squares fits to the data.

Both short and long-term dynamics of gut microbiota contribute to overall turnover in gut bacterial species. To directly investigate the dynamics of gut microbiota composition, we next calculated the distribution of residence (*t*_*res*_) and return times (*t*_*ret*_) for individual OTUs. Following previous macroecological analyses^9, 30, 31^, we defined residence times as time intervals between the emergence and subsequent disappearance of corresponding OTUs; analogously, return times were defined as the intervals between disappearance and reemergence of OTUs. Again, we observed residence patterns very similar to those previously described in diverse ecological communities^9, 30, 31^ (Supplementary Fig. 4b). Specifically, the distributions of *t*_*res*_ and *t*_*ret*_ were described well by power laws (Equation 3) with exponential tails resulting from the finite length of the analyzed time series (Fig. 3, Supplementary Fig. 9a,b, Supplementary Fig. 10).

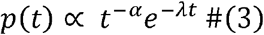

**Fig. 3.**
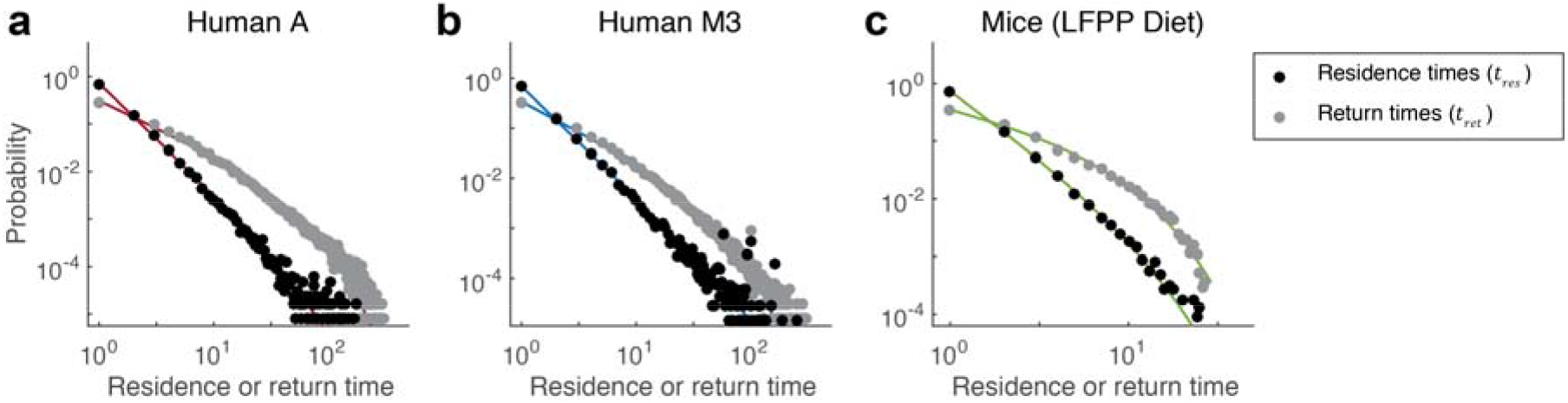
Residence and return times of gut microbiota. **a-c,** Residence (*t*_*res*_) and return times (*t*_*ret*_) were defined as the number of consecutive time points during which an OTU was detected at any abundance in the community or absent from the community respectively. Probability distributions for *t*_*res*_ and *t*_*ret*_ follow power laws of the form *p*(*t*) ∝ *t*^−*α*^*e*^−*λt*^, with the exponential tail resulting from the finite length of each time series. Power law exponents are *α*_*res*_ = 2.3 ± 0.04, 2.2 ± 0.07, 2.2 ± 0.04 for residence times and *α*_*ret*_ = 1.1 ± 0.02, 1.2 ± 0.05, 1.2 ± 0.07, 1 for return times (mean ± s.d., humans A and M3 and LFPP mice respectively, Methods). Residence and return times are aggregated across the three individual mice on the LFPP diet. Solid lines indicate fits to the data using MLE.

The residence times distributions were similar within and between individual human and mouse gut microbiomes (*α*_*res*_ = 2.3 ± 0.05, *α*_*ret*_ = 1.2 ± 0.02, mean ± s.d. across humans, *α*_*res*_ = 2.2 ± 0.04, *α*_*ret*_ = 0.72 ± 0.03, across mice on the LFPP diet), suggesting that the processes governing the local emergence and disappearance of gut bacteria are likely to be independent of the specific host. The power-law with an exponential tail distribution of residence times may arise, even in isotropic environment, from the dynamics of births, deaths, and species migration patterns defined by the spatial structure of the ecosystem^30^. Notably, the power law exponents (~2) of the residence distribution revealed in our analysis are similar to the ones observed previously in macro ecology^30^.

Having characterized distributions of daily abundance changes and residence times, we next investigated the temporal variability of individual OTU abundances. One of the most general relationships in ecology that has been observed across hundreds of different biological communities is known as Taylor’s power law^32–35^, which connects a species’ average abundance to its temporal or spatial variance,

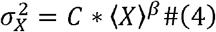

where *C* is a constant, ⟨*X*⟩ and 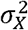 are the mean and variance of species abundances respectively, and *β* is a positive scaling exponent. For processes following simple Poissonian fluctuations, the parameter *β* = 1, while for processes with constant per capita growth variability^36^, *β* = 2. Values of *β* have been empirically observed to lie between 1 and 2 for the vast majority of investigated plant and animal species^37^. Interestingly, our analysis revealed that the temporal variability of gut microbiota also followed Taylor’s law (Fig. 4, Supplementary Fig. 11a,b), with exponents for human and mouse gut microbiomes generally consistent with values observed previously in other ecological communities^37^ (*β* = 1.7 ± 0.02 across humans, *β* = 1.49 ± 0.02 across LFPP mice). Notably, compositional effects of microbiota datasets did not explain the values of the observed Taylor law exponents (Supplementary Fig. 12). Dynamics consistent with Taylor’s law have been also observed in a recent short-term analysis of the healthy human vaginal microbiome^38^. It has been previously suggested that competitive interactions between species may result in Taylor’s law exponent in the range between 1 and 2 ^36^. Alternatively, a Taylor’s law with nontrivial exponents may arise due to stochastic demographics of population growth and decline^37^, presence of species subtypes each with a gamma-distributed abundances^39^, or due to a balance between tendencies of the species to aggregate and disperse ^35, 40^. In the future, it will be interesting to apply and compare these diverse theoretical models in the context of microbiota dynamics.

Although Taylor’s law described well the overall dynamics of gut microbiota, some specific OTUs clearly deviated from the general trend (Fig. 4). To determine whether their behavior reflected specific ecological perturbations, we identified all OTUs that exhibited significant and abrupt increases in abundance during previously documented periods of travel in human A and enteric infection in human B^7^ (Methods). Interestingly, these travel and infection-related OTUs corresponded to the outliers from Taylor’s law (Fig. 4a,b, blue circles), showing on average ~10-fold greater variance than expected based on the Taylor’s law trend (Supplementary Fig. 11a,c, Supplementary Table 1). Many of these OTUs were members of the Proteobacteria (OTU 13, family: Enterobacteriaceae, OTU 29, family: Pasteurellaceae, OTU 5771, family: Enterobacteriaceae in human A; OTU 13, family: Enterobacteriaceae in human B), which were associated with the microbiota perturbations^7^ (Supplementary Table 1). Moreover, other OTUs, primarily belonging to the Firmicutes, that exhibited abrupt changes in abundances (OTU 25, family: Peptostreptococcaceae in human A; OTU 95, family: Ruminococcaceae, OTU 110, family Ruminococcaceae in human B) also displayed higher than expected temporal variability (Fig. 4a,b, purple circles, Supplementary Fig. 11c, Supplementary Table 1). These results suggest that macroecological relationships can be used to identify and characterize specific microbial taxa that are likely involved in periods of dysbiosis and other context-specific environmental perturbations.

**Fig. 4.**
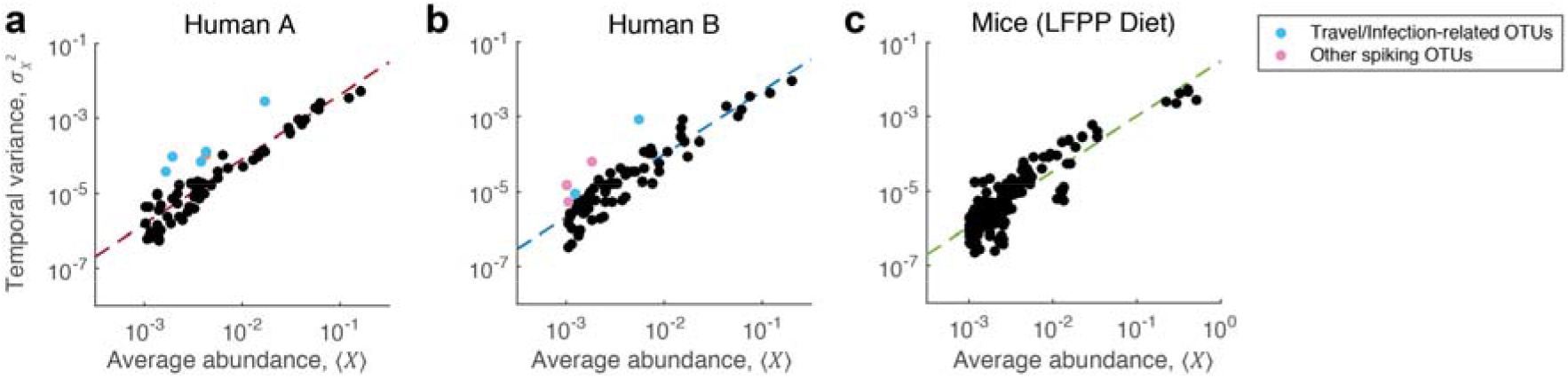
Taylor’s power law in the gut microbiome. Mean and temporal variance of OTU abundances follow Taylor’s power law of the form 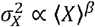, with *β* = 1.66±0.09, 1.60±0.08, 1.49±0.02 for humans A, B and LFPP mice respectively (mean ± s.d., see Methods). Each point corresponds to the average abundance and temporal variance of a single bacterial OTU. **a,b,** OTUs that exhibited temporary and abrupt increases in abundance are indicated as colored circles (Methods). Light blue circles indicate OTUs that exhibited significant increases in abundance specifically during periods of travel (human A) and enteric infection (human B). **c,** Data from each mouse on the LFPP diet are overlaid. Dashed lines indicate least-squares regression fits.

It is well established that the dynamics of diverse ecosystems are strongly affected by their environment^41^. Host dietary intake is a major environmental factor influencing gut bacterial abundances^8, 17, 42^ and disease phenotypes^43, 44^. Therefore, we next explored the effects of diet on the observed macroecological relationships describing gut microbiota dynamics. To that end, we used data from the study of Carmody et al.^8^, who investigated fecal bacterial abundances in individually-housed mice fed either a low-fat, plant polysaccharide-based (LFPP) diet, or a high-fat, high-sugar (HFHS) diet. Our analysis revealed that the short-term dynamics of gut microbiota were significantly affected by the diets. While the variability (standard deviation) of daily abundance changes declined rapidly with increasing abundance in the LFPP mice (Fig. 5a, green), it remained more homogeneous across OTU abundances in the HFHS mice (Fig. 5a, purple, regression slopes *r* = −0.17 ± 0.03 for the LFPP diet, −0.08 ± 0.02 for the HFHS diet, Z-test of regression coefficients p= 2.0e-5). As we describe below, the relatively smaller short-term variability of highly abundant species on the LFPP diet likely reflects more stable niches for bacteria (Bacteroidetes) that may catabolize dietary fibers. On the other hand, relatively higher fluctuations of lowly abundant bacteria on this diet may be induced by cross-feeding on catabolic products of highly abundant species. The observed dependence of short-term variability on species abundances is much weaker on the HFHS diet, which may result from a general loss of niche diversity due to the substantially reduced nutrient complexity of that diet.

**Fig. 5.**
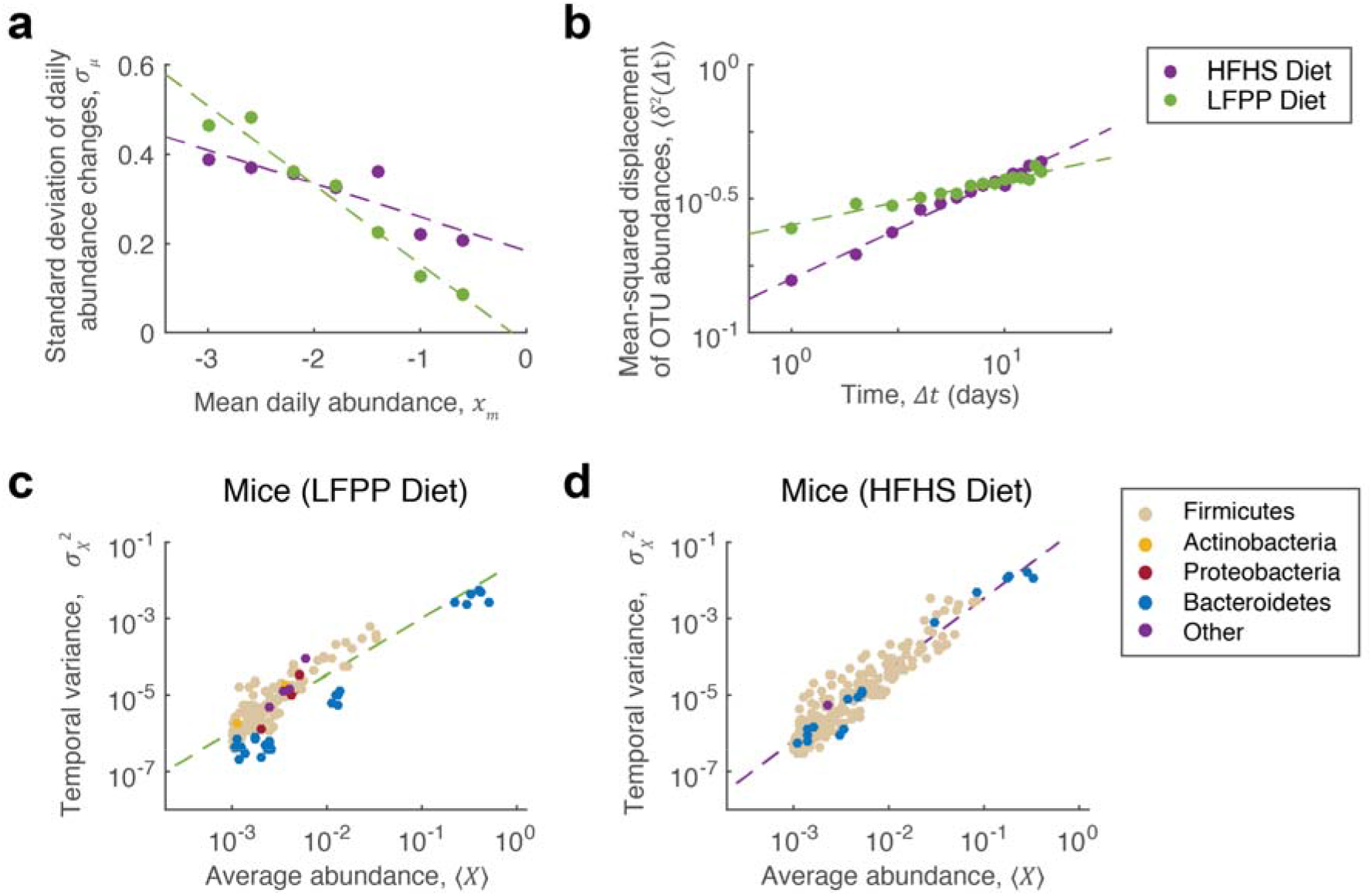
Dynamics of gut microbiota in mice fed different diets. **a,** OTUs in mice fed a low-fat plant-polysaccharide-based (LFPP) diet show a stronger dependence of the variability in daily abundance changes (*σ*_*μ*_) on mean daily abundance (*x*_*m*_) compared to those fed a high-fat high-sugar (HFHS) diet (regression slopes *r* = −0.17± 0.03, *r* = −0.08± 0.02; mean± s.d., LFPP and HFHS mice respectively). Data are aggregated across the three mice on each diet with dashed lines indicating least-squares regression fits. **b,** OTU abundances in the LFPP mice exhibit reduced long-term abundance drift compared to those in the HFHS mice (*H* = 0.08 ± 0.02, *H* = 0.19 ± 0.02). **c,d,** Taylor’s law analysis shows differences in overall scaling of average OTU abundance versus temporal variance on each diet (*β* = 1.49 ± 0.02, *β* = 1.86 ± 0.07), driven by the temporal behavior of the Bacteroidetes in the LFPP mice (blue circles). Plots correspond to data combined from the three mice on each diet. Dashed lines indicate least-squares regression performed on the combined data.

In other ecological communities, smaller short-term fluctuations of species abundances do not necessarily lead to increased long-term ecological stability and vice versa^45, 46^. Thus, in addition to short-term fluctuations, we also investigated how different diets affected the long-term drift of gut microbiota. Hurst exponents were significantly larger in the HFHS mice, indicating substantially faster long-term drift of bacterial abundances on this diet (Fig. 5b, Supplementary Fig. 7b, *H* = 0.19 ± 0.02 for the HFHS diet, 0.08 ± 0.02 for the LFPP diet, Z-test p<1e-10). We note that short-term fluctuations in abundances was somewhat higher on the LFPP diet compared to the HFHS diet, which is reflected in a higher Y-axis intercept for the diffusion on the LFPP diet (Fig. 5b); the higher intercept is due to larger short-term fluctuations of numerous low abundant bacteria on the LFPP diet compared to the HFHS diet (Fig. 5a). Despite the intercept differences, the observed diffusion trend continues over long time scales (>100 days, based on the data in Fig. 2). Therefore, the long-term drift of microbiota abundances is likely to primarily depend on the difference in the respective Hurst exponents.

Previous studies have demonstrated diet-induced compositional shifts of gut microbiota^8, 17, 42^ and a reduced gut bacterial diversity in Western populations attributed in part to altered dietary habits^47–49^. Our analysis shows that different diets not only affect the composition of gut microbiota, but also significantly change their long-term dynamics. The reduced long-term stability on the HFHS diet may result from a higher degree of neutral drift and increased inter-species competition associated with a more homogeneous nutrient environment^50^. In addition, we found that while the abundance drift of the Bacteroidetes and Firmicutes, two major phyla in the mouse gut, were relatively similar on the HFHS diet (*H* = 0.18 ± 0.1 for Bacteroidetes, *H* = 0.18 ± 0.03 for Firmicutes), the Bacteroidetes exhibited significantly reduced drift on the LFPP diet as compared to the Firmicutes (*H* = 0.03 ± 0.06, *H* = 0.09 ± 0.02 Z-test p=3e-8). This suggests that while the LFPP diet decreased the long-term abundance drift of all taxa, the stability of the Bacteroidetes was particularly affected by this diet (see below).

Different diets may not only change overall gut microbiota dynamics, but also alter the temporal variability of individual taxa relative to the rest of the community. To understand taxa-specific changes, we examined Taylor’s law in mice on the LFPP and HFHS diets (Fig. 5c,d). Interestingly, it has been previously demonstrated that the Taylor’s law exponent may significantly depend on the environment, at least for some species^51–53^. Our analysis of the gut microbiota dynamics on the different diets is consistent with these observations. Specifically, we found that power law exponents were significantly different between the two diets (*β* = 1.49 ± 0.02 for the LFPP diet, *β* = 1.86 ± 0.07 for the HFHS diet, Z-test p=1.5e-6). The temporal fluctuations of the Bacteroidetes (Fig. 5c,d, blue circles) exhibited significantly lower variability given their abundances on the LFPP diet, but not on the HFHS diet (hypergeometric test, p=2.4e-4, Supplementary Table 2, Methods). Moreover, we observed significantly lower Taylor’s law exponents on the LFPP diet specifically for the Bacteroidetes (*β* = 1.66 ± 0.06 on the LFPP diet, *β* = 1.95 ± 0.03 HFHS, Z-test p = 0.0023), but not for all other bacterial taxa (*β* = 1.84 ± 0.2 for the LFPP diet, *β* = 1.86 ± 0.07 for the HFHS diet; Z-test p = 0.39). Furthermore, the highly abundant Bacteroidetes and their lower temporal variability on the LFPP diet were primarily responsible for the relatively smaller short-term fluctuations of highly abundant bacteria on this diet (Fig. 5a). Bacteroidetes are known to metabolize a wide range of dietary fibers present in the LFPP diet^55–57^ and are significantly lost during multigenerational propagation of mice on a low-fiber diet^48^. This suggests that specific members of the Bacteroidetes (OTU 118, OTU 237, OTU 364, family: Porphyromonadaceae, Supplementary Table 2) may exhibit both lower temporal variability and abundance drift by directly exploiting stable niches that are present on the LFPP diet and likely lost on the HFHS diet. These results demonstrate that macroecological analyses can identify specific taxa whose temporal dynamics are altered between different diets.

Our study demonstrates that, in spite of an amazing interaction and organizational complexity, the dynamics of gut microbiota can be described by multiple robust quantitative relationships. These scaling laws characterize both short- and long-term microbiota dynamics, and are usually observed across many orders of magnitude in time and bacterial abundances. We furthermore show that these relationships are unlikely to arise due to technical noise, compositional nature of microbiome datasets, and effects associated with species aggregation. Despite the difference of more than six orders of magnitude in the relevant spatial and interaction scales, the statistical relationships described in our study are strikingly similar to those observed previously in many diverse macroecological systems. This similarity suggests that the temporal processes in both macroscopic and microbial communities may be governed by a universal set of underlying mechanisms and principles.

We anticipate that the quantitative statistical framework developed in macroecology^4, 58, 59^ will be important for analyzing microbiota dynamics. As the observed statistical relationships describe different aspects of community dynamics, an important goal for the future studies will be to unify these observations into an integrated view of microbiota ecology, which also takes into account the spatial and environmental dimensions. Moreover, the ability to easily perturb microbiota composition and environment, add and remove particular species, as well as monitor species abundances at high temporal and spatial resolution, suggests an exciting opportunity to use microbiome as a convenient model system to explore general ecological relationships.

We also envision that a quantitative ecological framework will be important for understanding how host-specific and environmental factors influence the dynamics of gut and other health-related microbiota. Our results suggest that the observed macroecological relationships can be used to identify both global dynamical changes and also specific taxa whose abnormal temporal behavior may serve as biomarkers for periods of illness and clinically relevant perturbations^61, 62^. Therefore, it will be important to investigate how the quantitative macroecological relationships revealed in our study vary across large and densely-sampled human cohorts^63–65^.s

## Methods

### 16S rRNA Sequence Analysis

Raw 16S rRNA sequencing data for humans A and B was obtained from the European Nucleotide Archive (accession number: PRJEB6518^7^). Raw sequencing data from humans M3, F4 and mice was obtained from the MG-RAST database^66^ (4457768.3-4459735.3 for humans; 4597621.3-4599933.3 for mice). Sequences were analyzed with USEARCH 8.1^67^ using an open clustering approach. For studies including unfiltered sequencing reads, filtering was performed using the -fastq_filtecommand with expected errors of 2. All reads were then truncated to 100bp, with shorter reads discarded. Following a conventional approach, reads were de-replicated and clustered at 97% sequence similarity using the – cluster_otus command to generate OTUs with a minimum of 2 sequences. Sequences were then assigned to OTUs using the -usearch_global command, resulting in OTU tables for each study. Taxonomic assignments were made to OTUs using the RDP classifier^68^. Sequencing reads from each sample were then rarefied to a depth of 25K, 17K and 25K for the two human studies (A/B, M3/F4) and one mouse study respectively using Qiime 1.8^69^.

### OTU Inclusion Criteria

To control for technical factors such as sample preparation and sequencing noise, analysis was restricted to OTUs passing two sets of criteria. First, OTUs were required to be present in over half of the samples within respective subjects. Second, OTUs were required to have a mean relative abundance > 1e-3 over the time series. The abundance cutoff corresponded to a mean of 25 (A, B, LFPP/HFHS mice) and 17 (M3 and F4) reads over respective sampling periods. The final analysis of human individuals included ~75 OTUs comprising ~90% of the reads assigned to an OTU in any given sample. For mice, these criteria resulted in the inclusion of ~70 OTUs in the HFHS diet and ~55 OTUs in the LFPP diet, comprising ~90% of reads assigned to an OTU in a given sample. Because the HFHS mice initially received a LFPP diet, the analysis of these mice began 5 days after the diet shift. For the calculation of residence and return times, different criteria were imposed (see below), as these analyses would be biased by a prevalence cutoff and were more robust to noise in OTU abundance levels.

### Daily abundance changes

Daily abundance changes were defined as *μ*_*k*_(*t*) = log (*X*_*k*_(*t* + 1)/*X*_*k*_(*t*)), where *X*_*k*_(*t*) is the relative abundance of a given OTU *k* on day *t*. Distributions reflect community averages, with abundance changes calculated for each OTU across all time points and aggregated over all OTUs. To estimate the variability of the distribution of daily abundance changes within human subjects, each time series was divided into six consecutive time frames of equal length (estimates were insensitive to this number). Within each time frame, daily abundance changes were calculated and maximum-likelihood estimation (MLE) was used to fit the Laplace distribution exponent, with the mean and standard deviation of these values reported in the main text. For the mouse study, standard deviations reflected variability across the three individual mice on each diet. Mean daily abundances *x*_*m*_ were defined as the mean of consecutive log OTU abundances, 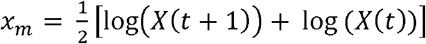. To estimate the variability in daily abundance changes as a function of abundance, abundance changes were binned by values of *x*_*m*_ using a bin size of 0.4 and standard deviations *σ*_*μ*_ were then calculated on the binned abundance changes. For diet comparisons, daily abundance changes were aggregated across the three mice on each diet. Abundance changes and mean daily abundances were calculated using the base ten logarithm in all figures, with the natural log used for parameter estimation.

### Hurst exponents

The mean-squared-displacement (MSD) of log OTU abundances was estimated as:

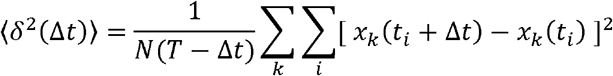

where the angled brackets denote a community average (over time and OTUs). Here, *x*_*k*_(*t*_*i*_) is the log relative abundance of OTU *k* at time *t*_*i*_, *N* is the total number of OTUs and *T* is the total length of the time series. A maximum time lag of 100 and 15 days were chosen for human and mice subjects respectively due to the finite length of each time series. Hurst exponents were then calculated by regressing ⟨*δ*^2^(Δ*t*)⟩ against Δ*t* on log-transformed axes. To estimate the variability of Hurst exponents within human subjects, time series were divided into six equal-length time frames as was done for daily abundance changes calculations. Hurst exponents for individual OTUs were estimated in a similar fashion but with displacements restricted to time averages. For diet comparisons, Hurst exponents were additionally averaged over mice within each diet:

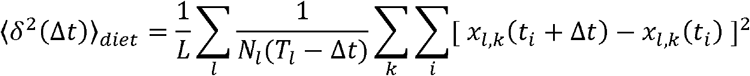

where the outermost summation is over individual mice (*L* =3) on each diet.

### Residence and return times

Residence times (*t*_*res*,*k*_) of an OTU *k* corresponded to the number of consecutive time points between its appearance (*T*_*a*,*k*_) and disappearance (*T*_*d*,*k*_) in the community (*t*_*res*,*k*_ = *T*_*d*,*k*_ − *T*_*a*,*k*_). Here, *T*_*a*,*k*_ is any time point at which the OTU was detected at a finite read count with no reads detected on the previous collection date, and *T*_*d*,*k*_ is the next time point at which reads were no longer detected. Return times (*t*_*ret*_) were similarly defined as the number of consecutive time points between local disappearance (*T*_*d*,*k*_) and reappearance (*T*_*a*,*k*_) in the community (*t*_*ret*,*k*_ = *T*_*a*,*k*_ − *T*_*d*,*k*_). Only intervals that fell entirely within the time frame of the study were included. A series of alternative criteria were also considered to ensure robustness of distributions. 1) To ensure results were not biased by detection sensitivity of sequencing, distributions were calculated for data subsampled to various sequencing depths (down to 1,000 reads per sample). 2) To account for false negatives in read detections, single read counts of zero, interrupting a run of consecutive nonzero abundances were neglected. That is, an OTU with zero reads at time was considered to be present in the community if that OTU was also present at times and *t* − 1 and *t* + 1. 3) To control for false positives, single read abundances were neglected and treated as a zero count. Results were qualitatively insensitive to both sampling depth and the alternative read detection criteria. To estimate variability of distribution parameters within human individuals, OTUs were randomized into six equal-sized groups. Residence and return times were calculated within each group and exponents were then fitted using MLE, with means and standard deviations reported in the main text. Within diets, means and standard deviations were calculated across individual mice.

### Taylor’s power law

The mean abundance ⟨*X*_*k*_⟩ and variance 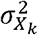 for each OTU *k* was calculated over the time series. Taylor’s exponents were obtained by performing linear regression of the log-transformed mean and variance across OTUs in each subject. To estimate variability of exponents within subjects, time series were divided into six consecutive time frames as described before. Spiking OTUs were defined as those whose abundance on any single day was greater than the average abundance over all other days by over 25-fold. Travel-related and infection-related OTUs in humans A and B were identified as those whose abundances spiked over 25-fold during the documented time periods^7^. For mice, Taylor’s law outliers were identified using a likelihood-based approach. Briefly, linear regression on the log-transformed means and variances were performed on all but a single OTU *k*. The probability of observing the left out OTU *k* was assigned using a Gaussian likelihood function based on estimated residuals. All OTUs with probability less than *α* = 0.025 were taken to be outliers. For diet comparisons, means and variances were aggregated across individual mice within diets groups.

### Simulations of sampling error associated with finite sequencing depth

Read counts were simulated using a multinomial distribution with parameters 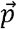 and *N*, where 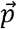 was estimated empirically from the set of average relative OTU abundances from humans A, B, M3 and F4 and *N* was taken to be the total sequencing depth used in each study (25,000 reads per sample for humans A and B; 17,000 reads per sample for humans M3 and F4). Simulations were performed multiple times to generate sample OTU trajectories in each human that solely reflected sampling error. To account for sporadic sequencing read dropouts, zero counts were introduced into simulations of each OTU to match the empirical frequency of zero counts observed in the real data.

### Simulations of microecological dynamics across a range of community diversities

Initial (*t* = *t*_0_) abundances of N=65 species were generated using power law distributions. The power law exponents were selected to generate a range of community diversities, quantified by the effective number of species (ENS = *e*^*H*^, where H is the Shannon diversity). For the simulated Gaussian daily abundance changes distribution (Supplementary Figure 3), Ornstein-Uhlenbeck process was used to generate 1,000 consecutive time points. For the Taylor’s law simulations (Supplementary Figure 12), we performed simulations identical to that of Kilpatrick and Ives^36^. Specifically, abundances of the 65 non-interacting species were propagated across 300 time points, matching the number of OTUs and samples in Human A. Both simulations were performed using absolute abundances, and the scaling macroecological relationships were then calculated using either absolute or relative abundances.

### Statistics

All statistical analysis was performed using custom scripts written in MATLAB (https://www.mathworks.com). Comparisons of various exponents between mouse diet groups were performed by first calculating the relevant coefficient and associated standard error of combined data across the three mice in each diet group. Z-tests were then performed comparing the two coefficients associated with each diet group assuming normality of standard errors. Reported p-values refer to one-sided tests.

## Data availability

All sequencing data used in this study can be downloaded from the ENA (https://www.ebi.ac.uk/ena/data/view/PRJEB6518 for humans A and B) and MG-RAST databases (https://www.mg-rast.org/linkin.cgi?project=mgp93 for humans M3 and F4; https://www.mg-rast.org/linkin.cgi?project=mgp11172 for mice). These data were used to generate all figures in the main text and supplement with the exception of Supplementary Figure 4.

## Code availability

All MATLAB scripts used to perform data analysis and generate figures will be available on GitHub at the time of publication.

## Author contributions

D.V. conceived the study. B.W.J., R.U.S., and K.T. performed data analysis. P.D.D. contributed to data interpretation. All authors wrote the manuscript.

## Materials and correspondence

Correspondence and requests for materials should be addressed to D.V.

